# The essential genome of *Xanthomonas citri*

**DOI:** 10.1101/2023.08.03.551896

**Authors:** Xiaolan Wang, Manying Wu, Yifei Ge, Weiwei Lv, Chaoying Liu, Xiaojun Ding, Yu Zhang, Jihua Wang, Yunzeng Zhang, Lei Li, Xiaofeng Zhou

## Abstract

Citrus canker, caused by the bacterium *Xanthomonas citri* subsp. *citri*, is a devastating disease with significant economic implications for the citrus industry worldwide. Understanding the molecular basis of *Xanthomonas* cell cycle and identifying therapeutic targets is crucial for effective disease management. In this study, we employed hyper-saturated transposon mutagenesis combined with high-throughput sequencing to determine the essential features of the *Xanthomonas citri* genome at ∼7-bp resolution. Our analysis revealed 525 essential genes, 181 high fitness cost genes, 7 small non-coding RNAs, 25 transfer RNAs, 4 ribosomal RNAs, and the origin of replication. Notably, the use of a newly designed Tn5 transposon with an outward pointing *lac* promoter significantly reduced false positives caused by polar effects associated with conventional transposons. Functional enrichment analysis showed that essential genes were significantly enriched in processes related to ribosome biogenesis, energy production and conversion, and membrane metabolism. Interestingly, the distribution of essential genes in *X. citri* showed similarities to that of the model organism *E. coli*, suggesting a conserved mode of genome organization that influences transposon accessibility. Our comprehensive analysis provides valuable target genes for potential therapeutic interventions against citrus canker and other related plant diseases.

## Introduction

*Xanthomonas citri* subsp. *citri* (Xcc) is recognized as the phytopathogen responsible for causing citrus canker, a disease characterized by erumpent pustules on the rind of fruits, leaves, branches, and seedlings. This highly pathogenic bacterium has a wide host range and can easily spread through rain, wind, and mechanical tools used in citrus plantations. It has been reported in various regions including Asia, South America, Oceania, and Africa, which are predominantly tropical or subtropical areas (Martins et al., 2020). Citrus canker poses a persistent threat to citrus yields and quality, leading to significant economic losses for growers. Despite the implementation of various disease management strategies such as the elimination of infected trees, the use of chemical compounds, and innovative approaches like generating transgenic trees and adopting biological control, these methods have proven to be inefficient, impractical, or require further exploration (Martins et al., 2020). Therefore, it is crucial to investigate the essential biological pathways that are vital for the survival of Xcc, as this knowledge can provide theoretical targets for the development of alternative preventive measures.

Essential genes are defined as genes that are required for the survival of an organism under specific environmental conditions (Luo et al., 2021). These genes and their corresponding products play crucial roles in the intricate metabolic network necessary for maintaining cellular life. Various technologies have been developed for the identification of essential genes (Peng et al., 2017). Among these, Tn-seq, a high-throughput method, has gained popularity for screening essential genes in bacteria (Cain et al., 2020; Nlebedim et al., 2021). Tn-seq combines saturation transposon-induced mutagenesis with next-generation sequencing, making it a robust, comprehensive, and highly efficient method that surpasses traditional techniques such as single-gene mutagenesis, experimental genome reduction, and antisense RNA (Juhas et al., 2011). The basic process of Tn-seq involves plasmid construction, generation of a transposon mutant library, identification of transposon insertion sites through high-throughput sequencing, and subsequent analysis of the sequencing results (van Opijnen et al., 2009, p.). The advent of Tn-seq has significantly accelerated the determination of essential genes on a genomic scale, particularly in prokaryotes, over the past decade (Luo et al., 2021).

The consideration of genetic context is crucial when assigning essentiality. Neglecting the genetic context can lead to common issues, such as the polar effect. The polar effect occurs when detrimental insertions happen in nonessential genes upstream of essential genes within the same operon, and it may have influenced previous studies’ determination of essentiality in operons (Morinière et al., 2021; Royet et al., 2018). However, in some cases, the impact of the polar effect may be minimal if the transposon used lacks a terminator that could disrupt the expression of downstream essential genes (van Opijnen et al., 2009; Wetmore et al., 2015). To mitigate the polar effect, a transposon that incorporates an active promoter are frequently used, which enables the generation of read-through transcripts. This approach helps to minimize the disruption of downstream essential genes caused by insertions within the same operon. (Christen et al., 2011; Goodall et al., 2018).

In this study, we aimed to identify the essential gene elements, including coding and non-coding genomic regions, that are crucial for the cellular growth of Xcc in a rich medium. To achieve this, we employed a hyper-saturated Tn5 transposon mutagenesis approach coupled with high-throughput sequencing (Tn-seq) with a resolution of nearly 7 base pairs (Figure 1). To address the issue of polar effects that can influence essential gene identification using conventional transposons, we engineered the Tn5 transposon to carry an outward pointing *lac* promoter and compared the transposon libraries generated by transposon with or without internal promotor (Figure 1). We conducted gene set enrichment analysis to analyze the functional genomics of the identified essential protein-coding genes. In addition, we conducted an analysis of the conservation of essential genes among the *Xanthomonas* genus and other related species within the Gamma-proteobacteria, which provided valuable insights into the comprehensive understanding of the organization and function of essential gene elements in *Xanthomonas* species. Ultimately, our findings may contribute to the identification of new drug targets for controlling this pathogenic bacterium.

**Figure 1.**
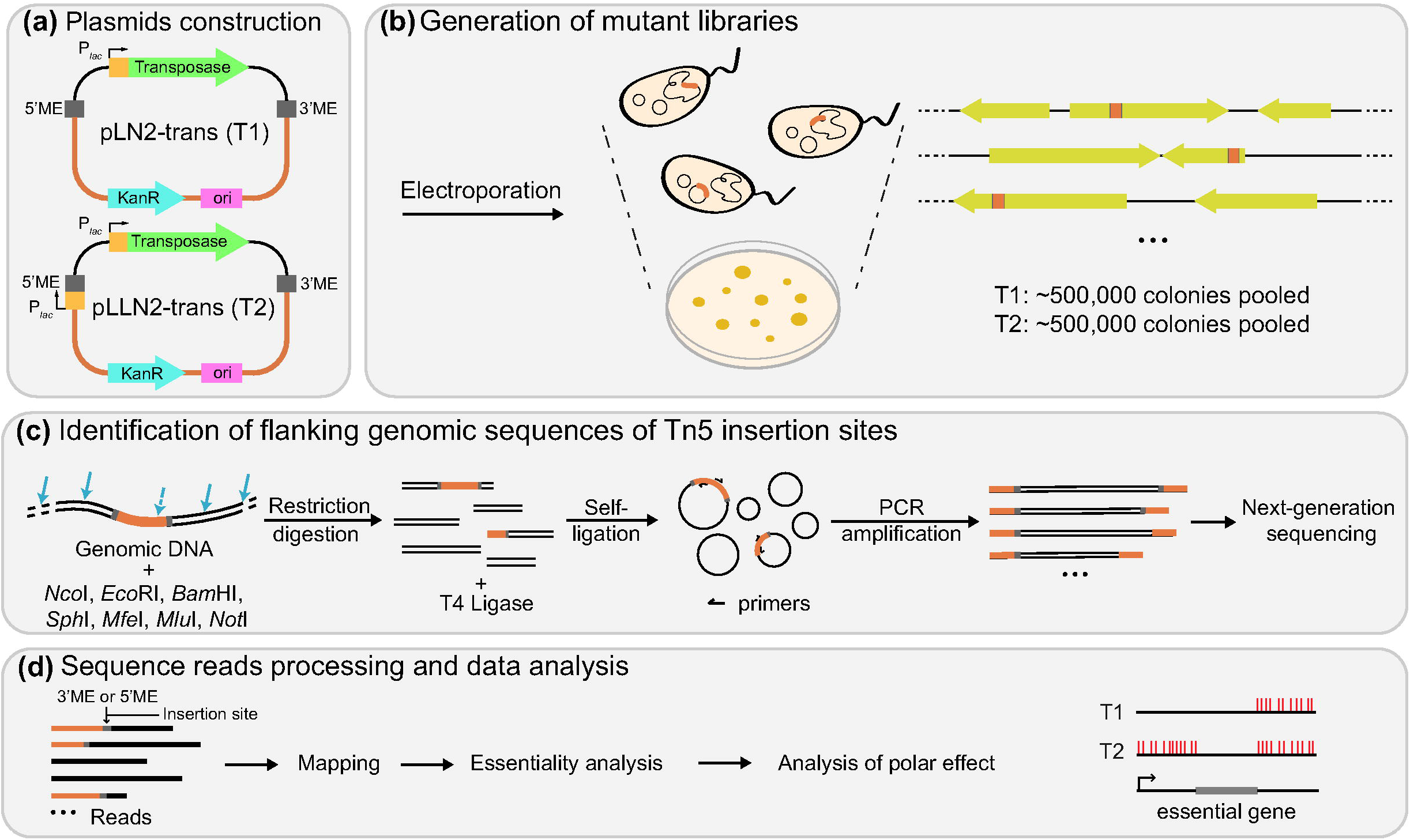
Steps in construction of hyper-saturated Tn5 mutant library for essential gene identification in Xcc. **(a)** Two plasmids used for creation of two Tn5 mutant libraries. The orange indicates the Tn5 transposon sequence. The pLLN2-trans contained an extra outward pointing *lac* promoter at one end of the Tn5 transposon. **(b)** Generation of mutant libraries by electroporating pLN2-trans and pLLN2-trans into Xcc CQ13 cells. Both T1 and T2 libraries contained ∼500,000 individual colonies. **(c)** Identification of flanking genomic sequences of Tn5 insertion sites. Genomic DNA extracted from each Tn5 library was digested by restriction enzymes followed by self-ligation. The ligation product was purified and subjected to library preparation for Illumina sequencing. **(d)** Sequence reads processing and data analysis. Sequencing reads containing 3’ or 5’ME sites with flanking sequences were filtered and mapped to CQ13 genome. The essentiality and polar effect analyses were then performed.

## Results

### Construction of highly saturated Tn5 mutant libraries in *Xanthomonas citri*

To construct a highly saturated Tn5 transposon insertion mutant library for the identification of essential genes in Xcc, we initially attempted to introduce pMCS2-transposase into Xcc CQ13 through electroporation. The pMCS2-transposase plasmid has been successfully used to generate a highly saturated Tn5 mutant library in *Caulobacter* (Christen et al., 2011). However, we encountered difficulties in obtaining a sufficient number of Tn5 transposon mutants using the pMCS2-transposase plasmid. This may be attributed to the low activity of the native promoter of *tnp*, which encodes a hyperactive form of the Tn5 transposase. To address this issue, we engineered the plasmid by replacing the native promoter of *tnp* with the *lac* promoter, which has been previously demonstrated to be constitutively active in *Xanthomonas*. The transformation of the modified plasmid, named pLN2, resulted in the generation of approximately 5 × 10^6^ viable colonies on nutrient agar plates. These colonies were subsequently pooled together to form the mutant library T1.

Next, we proceeded to construct a second plasmid called pLLN2, which incorporated an outward pointing *lac* promoter at one end of the Tn5 transposon. This modification aimed to introduce a new transcription initiation site downstream of the insertion, potentially mitigating the impact of the polar effect. By reducing the disruption of downstream gene expression in the same operon caused by insertions in upstream genes, the outward pointing *lac* promoter played a crucial role in minimizing the influence of polar effects. Similar to the previous procedure, we performed electroporation of pLLN2 into the Xcc CQ13 strain and successfully obtained approximately 5 × 10^6^ viable colonies. These colonies were subsequently pooled to form the mutant library T2.

To determine the transposon insertion sites, we employed multiple transposon non-cutting restriction enzymes to digest the isolated genomic DNA from both libraries. The digested fragments were then ligated using T4 ligase. The genomic regions flanking the transposon insertion sites were subsequently amplified in a reverse orientation using a pair of outward primers designed to bind specifically to the transposon sequence. We employed an Illumina Hi-seq system to obtain sequencing data from the two independent transposon libraries (Table 1). The raw sequencing data yielded 70,433,254 and 36,761,964 sequence reads for library T1 and T2, respectively. These reads were subjected to filtering based on the matching of each sequence read to the specific O-end of the transposon sequence. As a result, we obtained 1,905,270 and 1,830,884 high-quality sequence reads that could be mapped to the Xcc CQ13 genome for library T1 and T2, respectively. This allowed us to identify 383,416 and 455,413 unique insertion sites for library T1 and T2, respectively (Figure 2a). Furthermore, Pearson analysis demonstrated a strong correlation in transposable insertion abundance between library T1 and T2, indicating the reliability of our data (Figure 2b). Therefore, we combined the data from both libraries, resulting in a total of 726,909 unique insertion sites. The high density of unique insertion sites corresponds to an average of one insertion every 7.07 base pairs. An example of a genome locus is illustrated in Figure 2c.

**Table1:**
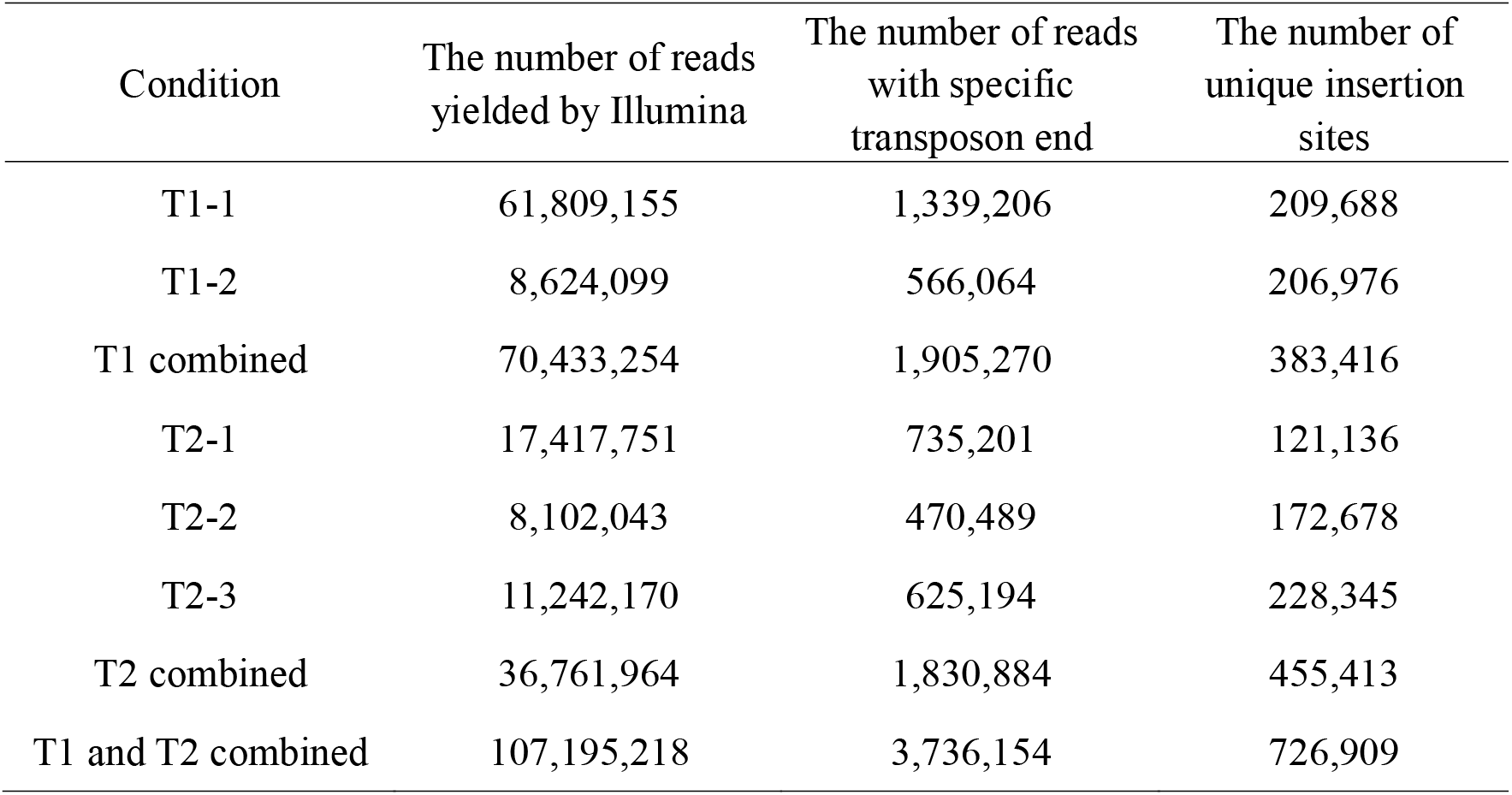
Parameters derived from *X. citri* CQ13 transposon insertion library.

**Figure 2.**
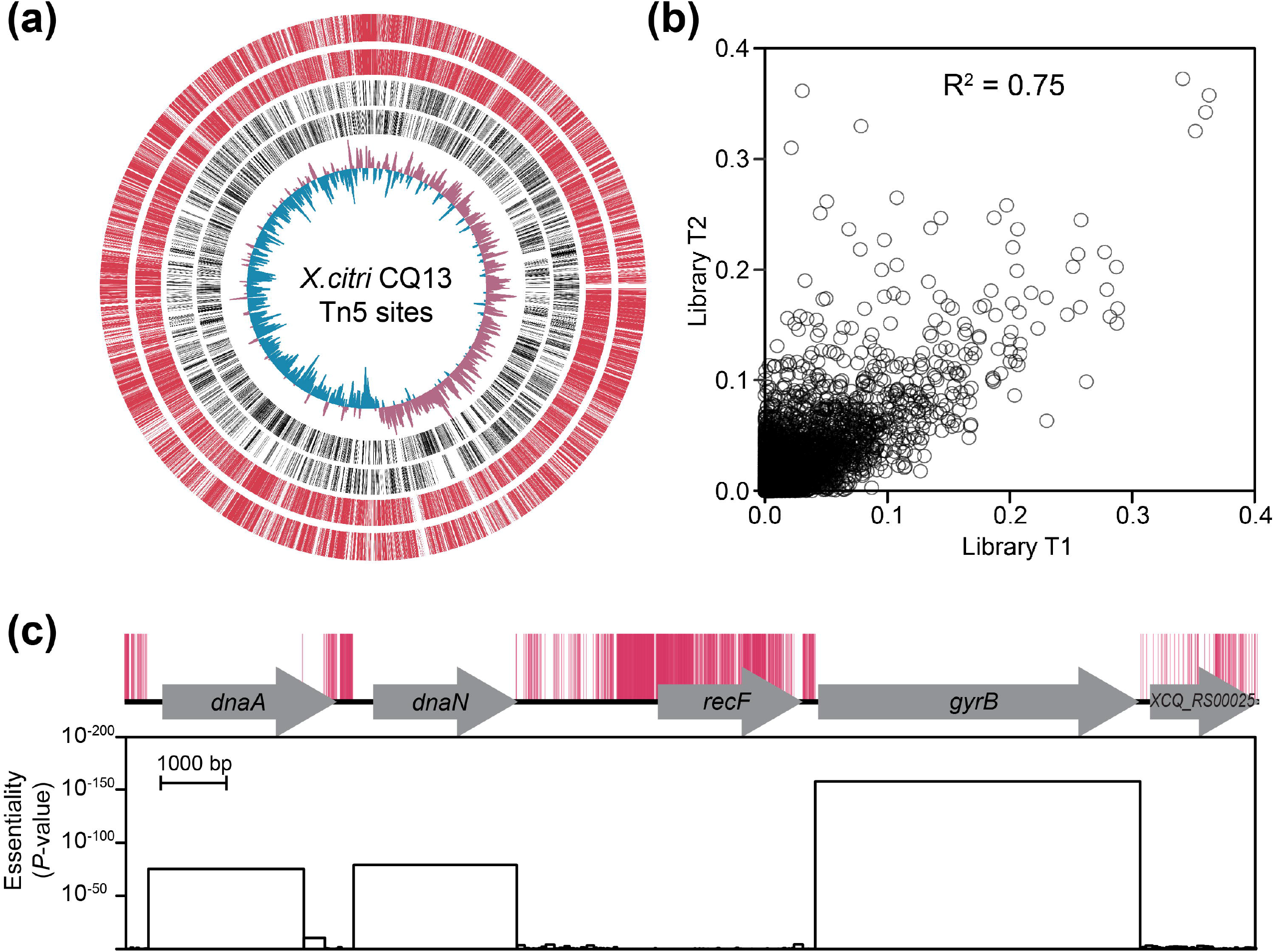
Complete essential genome atlas of *X. citri* CQ13. **(a)** Distribution of transposon insertion sites identified by Tn-Seq across the entire genome. The outer two red lines represent 383,416 and 455,413 unique transposon insertion sites identified from T1 and T2 libraries, respectively. The two adjacent black circles represent protein-coding genes on the forward and reverse strands, respectively. The innermost track represents the GC-skew, with positive values shown in maroon and negative values in blue. **(b)** Pearson correlation coefficients between the T1 and T2 libraries. **(c)** Locations of mapped transposon insertion sites (red marks) on the locus of the first five genes of the Xcc genome. For every non-disruptable genome region, a *P*-value for gene essentiality is calculated assuming uniform distributed Tn5 insertion frequency and neutral fitness costs.

### Essential and high-fitness cost gene elements

The assessment of gene essentiality using Tn-seq typically relies on the number of insertion sites within a specific gene or chromosome region. Conventionally, a gene with no insertion sites is classified as essential (Duong et al., 2018). In this study, we employed Tn5Gaps, a method provided by TRANSIT, to evaluate gene essentiality (Griffin et al., 2011). Instead of merely counting insertion sites, this approach defines essential open reading frames (ORFs) by detecting statistically significant gaps in transposon insertion coverage. Through this method, we identified 525 essential ORFs and 181 high-fitness cost ORFs on the chromosome (additional 7 essential ORFs on the two plasmids), which exhibited a substantial impact on bacterial fitness upon disruption by the Tn5 transposon (Supplementary Table 1).

We discovered that 270 essential ORFs did not have any insertions, indicating that their full-length ORFs were necessary for their function (Supplementary Table 2). An example of such an ORF is *gyrA* (Figure 3a). Among the 255 essential ORFs with Tn5 insertions, 75.8% of the insertions were concentrated within 5’- or 3’-terminal regions of inserted genes (Figure 3b). Within this group, 95 essential ORFs exhibited Tn5 insertions within their 3’ regions, enabling the identification of non-essential protein domains within these essential genes (Supplementary Table 3). For instance, the ATP-dependent Clp protease *clpP* gene displayed 43 Tn5 insertions within the last 115 base pairs of its 3’ region, which corresponded precisely to a ClpB_D2-small domain comprising a mixed alpha-beta structure. The disruption of the C-terminal D2-small domain of ClpP by Tn5 insertions may lead to the re-stabilization of essential protein substrates, rendering it dispensable for bacterial viability (Figure 3c). Additionally, we found 48 essential ORFs that only contained insertions at the 5’ region (Supplementary Table 4). For instance, the gene XCQ_RS13920, encoding an aspartate β-semialdehyde dehydrogenase (ASDase), had 33 Tn5 insertions near the originally annotated start codon, indicating a potential mis-annotation of the gene (Figure 3d). BLAST analysis also showed that the start codon of ASDase in other bacteria was identical to our corrected annotation in Xcc (Supplementary Figure 1). In total, we identified 27 mis-annotated genes in the CQ13 genome (Supplementary Table 5).

**Figure 3.**
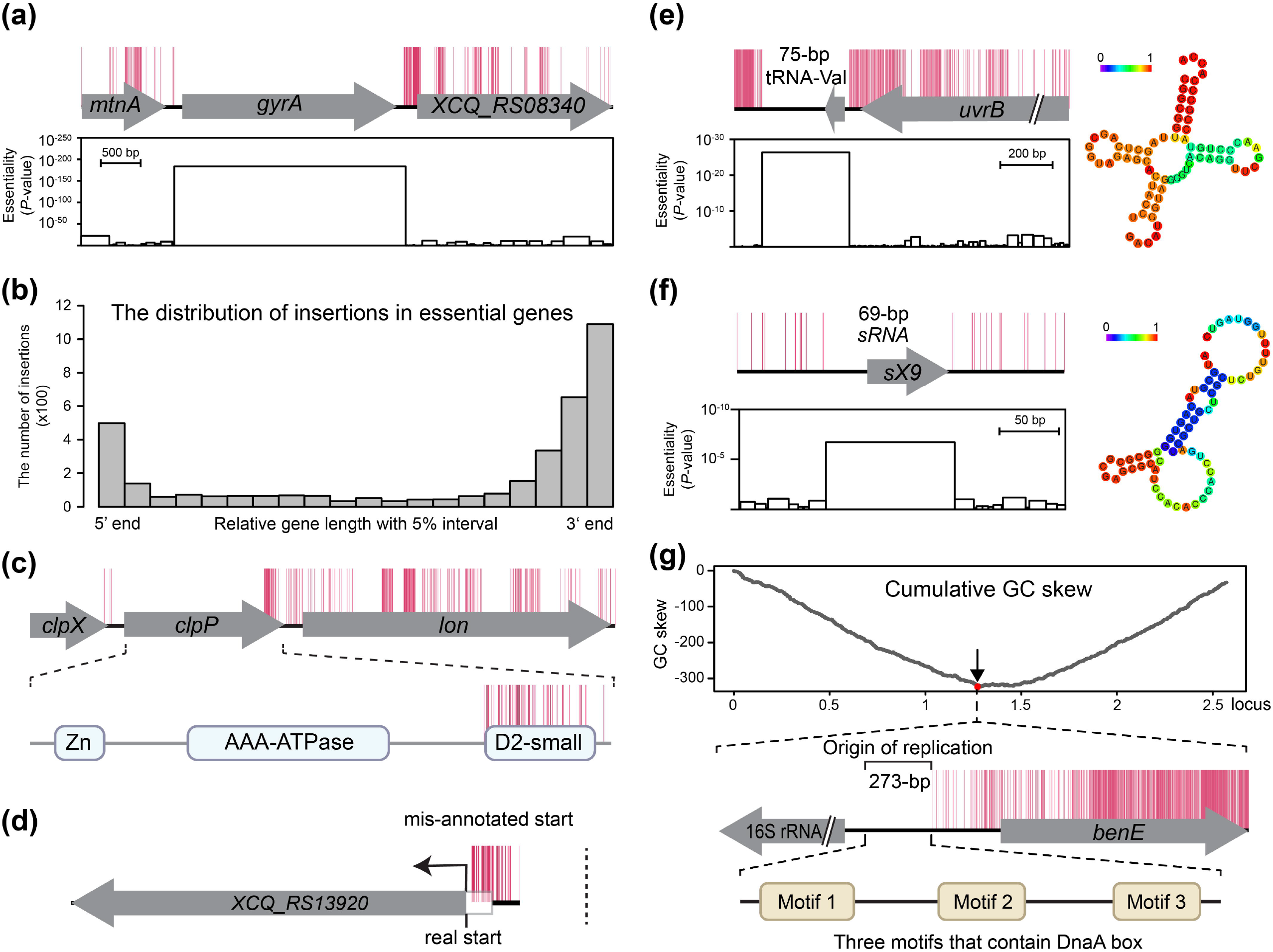
Identification of essential gene elements. **(a)** Essential gene *gyrA* showing no Tn5 transposon insertions across the entire ORF. **(b)** Distribution and enrichment of insertion sites for the 255 essential genes that allow transposon insertions at the 5’ or 3’ end. **(c)** The essential gene *clpP* had multiple transposon insertions within the 3’ end. This dispensable region encodes a D2-small domain that is non-essential for the function of ClpP. **(d)** Gene XCQ_RS13920 tolerates Tn5 insertions at the 5’ end of the mis-annotated ORF. The correct start site is 66 bp away from the original start site, and only 9 bp from the nearest transposon insertion. **(e)** A small non-disruptable segment containing an essential tRNA. **(f)** A 69-bp non-disruptable segment containing an essential sRNA. **(g)** Schematic representation of the transposon insertion pattern around the origin of replication. The top panel shows the cumulative GC skew across the whole genome. The minimum value of the cumulative GC skew across the entire genome was identified as the origin of replication. The zoomed-in section highlights the transposon insertion pattern and the predicted motifs, which are located at positions 4-53 bp, 100-149 bp, and 201-250 bp within the replication origin region.

We also investigated non-coding regions across the genome that did not exhibit any insertions. Within these regions, we identified 36 non-disruptable RNAs (Supplementary Table 6), including 7 small non-coding RNAs (ncRNAs), 25 transfer RNAs (tRNAs), and 4 ribosomal RNAs (rRNAs), as well as the origin of replication in the Xcc genome. For instance, the tRNA-Val, responsible for transporting valine, and a small ncRNA sX9 were found to be essential (Figure 3e and 3f). Additionally, we utilized GC cumulative skew analysis, which is particularly valuable in determining the location of replication origin and terminus (Bentley and Parkhill, 2004), and successfully identified the replication origin of the Xcc chromosome (Figure 3g). The replication origin was situated in an intergenic region spanning 5,038,400 to 5,038,600 base pairs, positioned between a 16S rRNA gene and a gene encoding a BenE family transporter, approximately 10 kilobases away from the chromosome replication initiator DnaA. This 273-base pair region contained three conserved putative AT-rich motifs that were identical to the DnaA box observed in Gamma-proteobacteria (Supplementary Figure 2).

### Comparison of T1 and T2 eliminates false positives identification of essential genes affected by polar effect

The identification of essential genes can be influenced by the polar effect of the inserted sequence, which may impact the expression or function of downstream genes within an operon, leading to false-positive results. To address this issue, we utilized T1 and T2 libraries with four types of Tn5 insertions. Importantly, only sense insertions in T2 allowed us to precisely identify false-positive essential genes within an operon containing other downstream bona fide essential genes (Figure 4a). To compare the distribution of insertions between T1 and T2 libraries, we calculated the ratio of insertions within a given gene to the total insertions in open reading frames (ORFs) across the genome for both libraries. The overall insertion patterns in each gene were found to be similar between T1 and T2, indicating that the two libraries were uniformly constructed (Figure 4b). However, the difference in this insertion ratio (T1-T2) revealed that miscounted essential genes occurred in the T1 library, suggesting that the polar effect was responsible for the false-positive identification of essential and high-fitness cost genes (Figure 4c).

**Figure 4.**
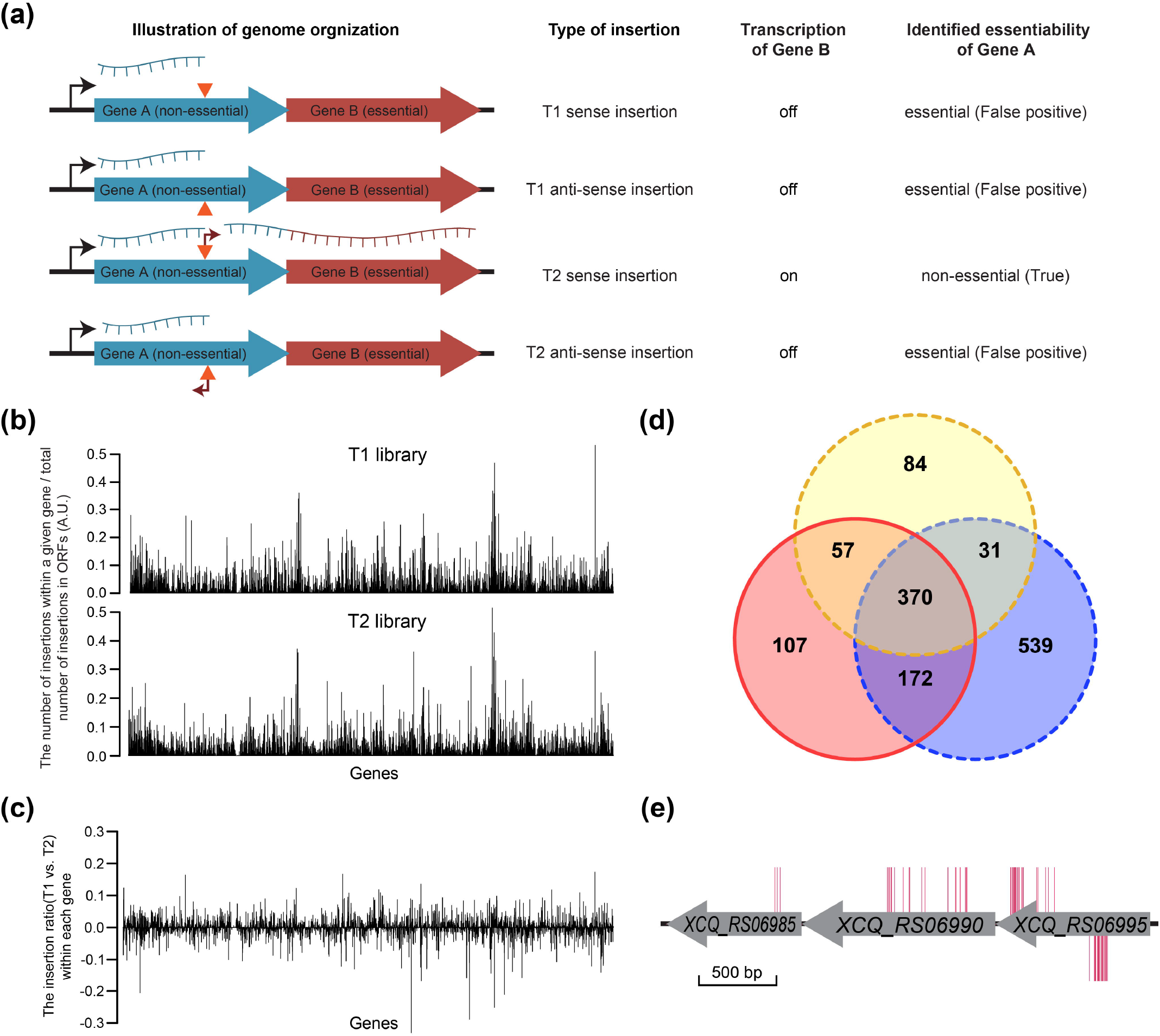
Comparison of T1 and T2 eliminates false positives of essential genes affected by polar effect. **(a)** Illustration of the pattern and orientation of Tn5 insertions in T1 and T2 libraries. Four types of Tn5 insertions were observed in both T1 and T2 libraries. Only sense insertions in T2 allowed precise localization of false-identified essential genes within an operon that contains other downstream bona fide essential genes. **(b)** Transposon insertion abundance in T1 and T2 libraries. The abundance was calculated as the number of transposon insertions in a given gene divided by the total number of transposons in T1 or T2. **(c)** Differences of transposon insertion abundance between T1 and T2 libraries. **(d)** The Venn diagram depicts the overlap and differences in essential and high-fitness cost genes identified in different libraries. The blue circle represents genes identified in the T1 library, and the yellow circle represents genes identified in the T2 library. The red circle represents genes identified in the integrated library of T1 and T2. **(e)** An example of the T2 library eliminating false positive of essential genes affected by polar effect. In a three-gene transcription unit (XCQ_RS06985 to XCQ_RS06995), only sense insertions in the T2 library occur on XCQ_RS06990, indicating that XCQ_RS06990 was false-positively identified as an essential gene in the T1 library. Red marks above the genes indicate the insertion patterns in the T2 library, while red marks below the genes represent the distribution of insertions in the T1 library.

The Tn5Gaps analysis revealed a total of 1112 and 542 essential genes, which included high-fitness genes, from the T1 and T2 libraries, respectively. Upon combining the libraries (T1 + T2), a set of 706 essential and high-fitness genes were identified (Figure 4d). Among the 539 essential genes that were exclusive to the T1 library, 32 of these genes were found in the upstream region of an identified essential gene within the same operon. The presence of these genes in close proximity to essential genes raised concerns that they might have been falsely identified as essential in the T1 library due to the polar effect caused by the disruption of downstream transcription within the same operon. However, to address this issue, a rigorous comparison between the T1 and T2 libraries was conducted. Notably, the results from the T2 library provided crucial information in distinguishing true essential genes from false positives due to polar effects. As an example, gene XCQ_RS06995 was consistently classified as non-essential since it had transposon insertions in both the T1 and T2 libraries (Figure 4e). Within the same operon, two downstream genes, XCQ_RS06990 and XCQ_RS06985, were initially identified as essential genes with no insertions detected in the T1 library. However, upon analyzing the distribution and orientation of transposon insertions in the T2 library, it was found that XCQ_RS06990 contained 18 insertions, all of which were inserted into the sense strand. These sense insertions enabled the transcription of downstream bona fide essential gene XCQ_RS06985, effectively ruling out the possibility of XCQ_RS06990 as an essential gene.

### The function of essential genome

To gain insights into the key functions of the essential gene set, we assigned these genes to functional categories or metabolic pathways using the COG database. The results of COG grouping revealed that approximately one third of the essential genes fell into three categories: 15% were associated with translation, ribosomal structure, and biogenesis; 9.3% were linked to energy production and conversion; and 7.6% were involved in cell membrane and envelope metabolism (Figure 5a). We also identified 62 essential genes that were categorized as having an unknown function (Figure 5a). To accurately assess the enrichment of COG categories within the essential gene set, we took into account their representation in the complete genome, thereby determining the degree of enrichment compared to their expected presence. Our results showed that five COG pathways, namely translation, ribosomal structure and biogenesis (J); nucleotide transport and metabolism (F); coenzyme transport and metabolism (H); energy production and conversion (C); and cell cycle control, cell division, chromosome partitioning (D), displayed significant enrichment (Log2 fold-enrichment close to or greater than 1) in the essential protein-coding gene set (Figure 5b). Four COG pathways, with Log2 fold-enrichment ranging from 0.47 to 0.11, demonstrated a slight enrichment, including intracellular trafficking, secretion, and vesicular transport (U); replication, recombination, and repair (L); cell wall/membrane/envelope biogenesis (M); and posttranslational modification, protein turnover, chaperones (O) (Figure 5b). Furthermore, we performed KEGG pathway enrichment analysis to obtain a more detailed clustering of essential genes. All the KEGG pathways depicted in Figure 5c showed a significant degree of enrichment within the set of essential genes. The majority of genes involved in aminoacyl-tRNA biosynthesis, cell cycle regulation, protein export, ribosome structure, and terpenoid backbone biosynthesis were deemed essential. Additionally, approximately half of the genes participating in DNA replication, peptidoglycan biosynthesis, and the citrate cycle were likewise indispensable for cellular viability of cells.

**Figure 5.**
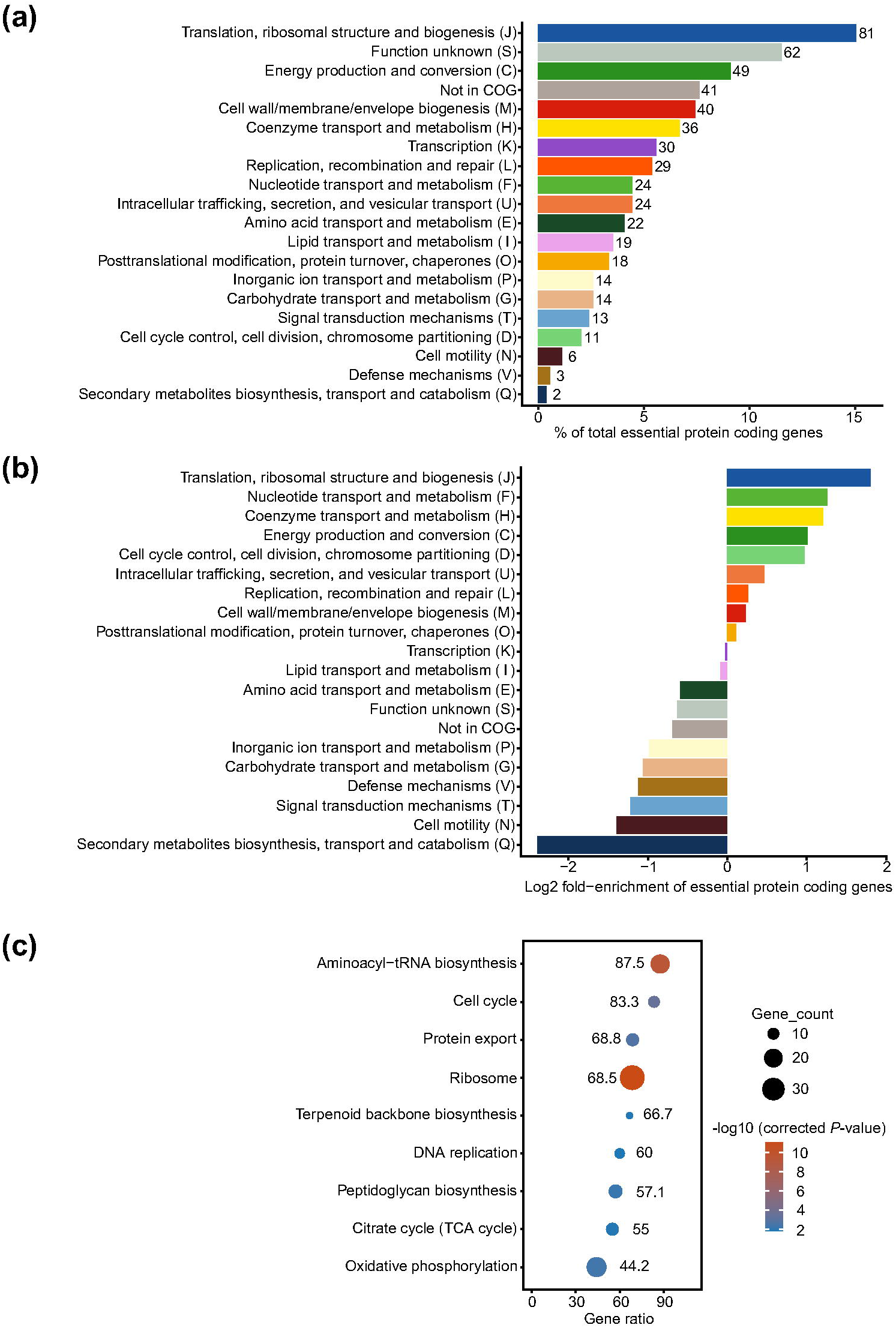
Clustering of Essential Genes based on Cluster of Orthologous Groups (COG) and KEGG Functional Categories. **(a)** The count of genes in each COG category within the essential gene set is depicted, with the corresponding numbers next to the bars representing the gene counts in each category. **(b)** Log-fold enrichment of COG categories, calculated as the ratio of the proportion of a given class in the essential gene set to its overall proportion in the genome. **(c)** Clustering of essential genes based on KEGG functional categories. Each dot represents a functional category. The size of dots represents the number of genes. The color indicates corrected *P*-values.

### Conservation and distribution of essential genes

To determine whether the essential genes identified from Xcc CQ13 were conserved, we searched for homologous sequences in the bacterial essential gene database of the DEG15 (Luo et al., 2021), which includes 66 bacterial essential gene sets. Out of the 525 essential protein-coding genes, 78.6% had putative homologues in the bacterial essential gene database, while 21.4% did not have any putative homologues.

Next, we focused our search within the genus *Xanthomonas* and other important model bacteria to identify specific essential genes from a phylogenetic perspective. We constructed a phylogenetic tree using the complete genomes of 15 bacteria from *Xanthomonas* and 3 bacteria from Gammaproteobacteria that were outside the genus *Xanthomonas* (Figure 6a). The majority of branches in our phylogenetic tree had a bootstrap support value greater than 80%, indicating a high degree of confidence in the depicted evolutionary relationships. Analyzing the conservation of the 525 essential protein-coding genes of Xcc CQ13 across the tree, we found that 454 out of the 525 essential genes were shared among all 15 bacteria from *Xanthomonas*. Among the remaining 71 genes, at least one absence was observed within the 14 *Xanthomonas* species, except for Xcc 306 (Figure 6a), which aligns with our previous study indicating that CQ13 serves as an alternative model strain compared to strain 306. (Wu et al., 2022). For the other three bacteria outside the *Xanthomonas* genus located at the bottom of the tree, only a small portion of the 525 essential genes were shared with Xcc CQ13.

**Figure 6.**
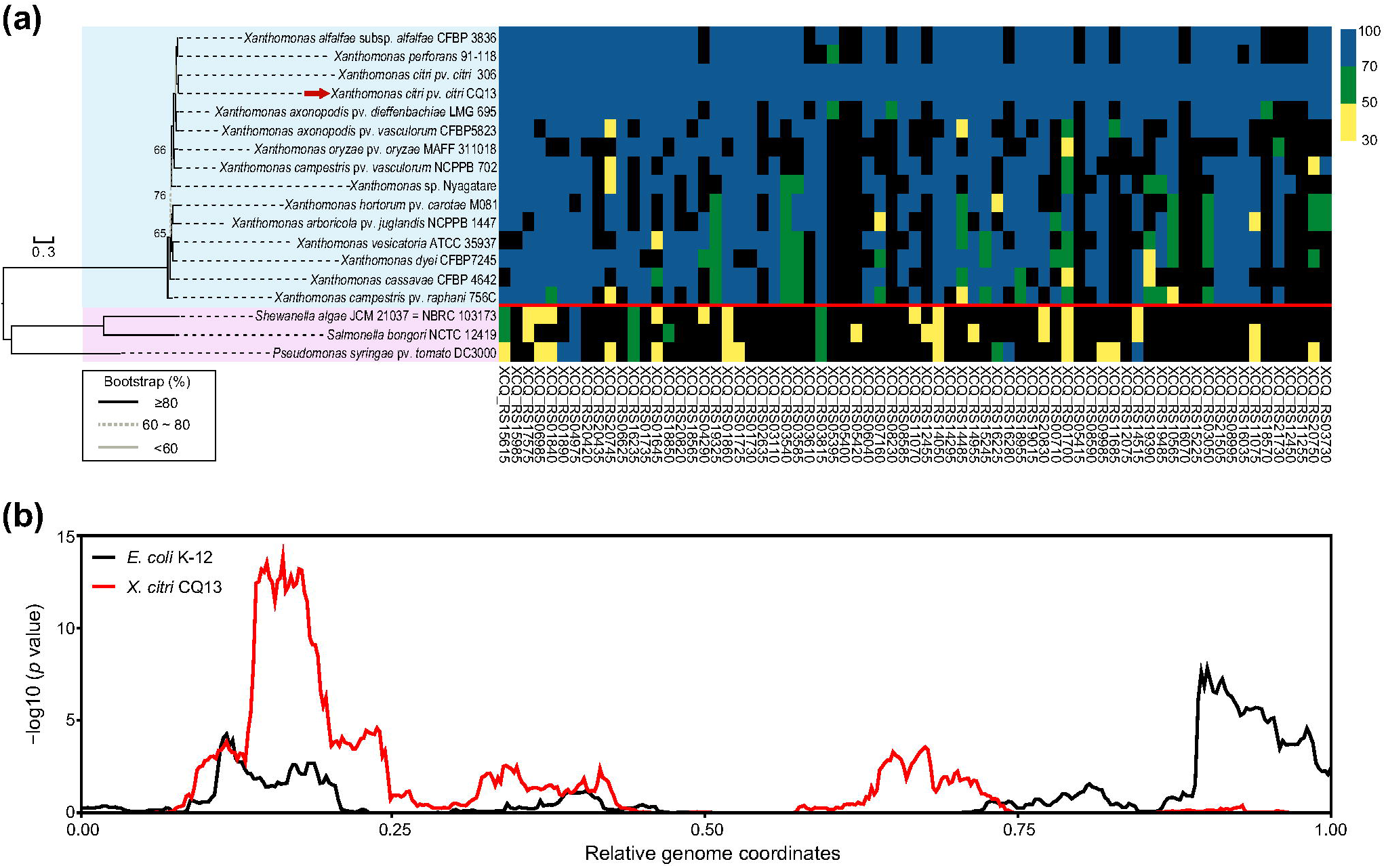
Conservation analysis of essential genes in Xcc CQ13. **(a)** Phylogenetic tree and conservation analysis based on the essential gene set in 15 *Xanthomonas* strains (blue) and 3 strains from other genera (pink). The CQ13 genome is indicated by a red arrow in the phylogeny. The taxonomy of *Xanthomonas* and other genera is separated by a red line. The gene conservation matrix displays the conservation level of essential genes across the analyzed strains. Blue boxes represent genes that were detected with BlastP using criteria of over 70% identity, green boxes indicate genes with identity ranging from 50% to 70%, and yellow boxes represent genes with identity ranging from 30% to 50%. Black boxes indicate genes with identity below 30%. Gene locus numbers are shown below the matrix. **(b)** Enrichment distribution of the essential gene set in the *X. citri* CQ13 genome (red) compared to the *E. coli* K-12 genome (black).

To examine the distribution and enrichment degree of essential genes along the Xcc genome, we calculated *P*-values to represent the degree of enrichment. In addition, we compared the distribution and enrichment of the essential gene set in Xcc with that of the model microorganism *E. coli*. In both Xcc and *E. coli*, essential genes were found to be unevenly distributed along the genome (Figure 6b). We defined the complete genome of Xcc from 0 to 1, with XCQ_RS00005 (*dnaA*) as the starting point and XCQ_RS21995 (*rpmH*) as the endpoint. The highest enrichment degree was observed at approximately 0.2 of the complete genome, while lower enrichment degrees were observed around 0.5 and 0.75 (Figure 6b). In *E. coli*, essential genes were mainly enriched in the either side of the origin (Goodall et al., 2018). To highlight the differences in essential gene distribution between the two bacteria, we overlapped the two distribution curves. Interestingly, the enrichment degrees of the two curves displayed a similar changing trend in the first half of their genome location, while the trend diverged in the second half of their genome location (Figure 6b). This finding suggests that the distribution of essential genes in both bacteria may reflect a conserved mode of genome organization that influences the transposon accessibility.

## Discussion

We utilized a combined approach of transposon mutagenesis and Illumina sequencing technology to identify genes essential for the survival of Xcc CQ13 in NB medium. Our results revealed that 11.9% (525) of genes on the Xcc chromosome are essential, which is in line with recent reports on other *Xanthomonas* species (Morinière et al., 2021; Pang et al., 2023). The 7-bp resolution of Tn5 insertions in our study allowed us to identify mis-annotations and critical domains in essential genes. Furthermore, we investigated the polar effect on partial genetic classification by comparing the insertion of transposons carrying or lacking a promoter on the transposon fragment. This enabled us to precisely rule out the false positives of essential genes located upstream of other essential genes within the same operon. Subsequently, we conducted a comprehensive analysis of the function, conservation, and distribution of the identified essential genes. Specifically, in the examination of genes related to energy production, we made a notable observation: most essential genes lacked complementary paralogs, indicating the presence of unique metabolic pathways that support the growth of Xcc in the medium. This finding suggests that these essential genes play crucial roles in specific metabolic processes essential for the bacterium’s survival and growth. Given the significance of essential genes in the functioning and survival of *Xanthomonas*, we recognize the importance of focusing on essential genes with unknown functions. Exploring these genes could provide valuable insights into the identification and understanding of vital pathways in *Xanthomonas*, shedding light on key processes that influence bacterial physiology and virulence.

In our analysis of transposon insertion sites, we observed an uneven distribution of Tn5 transposons across the entire genome of Xcc. Remarkably, when we combined the data of whole genome essential gene distribution, we found a significant enrichment of Tn5 insertions at the origin and termini of replication. Interestingly, this distribution pattern is in contrast to a previous report in the alpha-proteobacterium *Caulobacter crescentus*, where the origin and termini of replication were shown to be resistant to Tn5 insertions (Christen et al., 2011). The resistance to transposons in *Caulobacter* was attributed to the binding of numerous accessory factors for chromosome segregation and replication, which limited their accessibility to transposons. However, the opposite observation in Xcc could be explained by a study that demonstrated chromosome segregation in Xcc to be asymmetrical (Ucci et al., 2014). This asymmetry in chromosome segregation may alter the distribution of transposon insertion sites, leading to the significant enrichment of Tn5 insertions at the origin and termini of replication in Xcc.

During our investigation, we conducted a detailed analysis of the genome loci where essential genes were enriched (Figure 6b). Interestingly, we observed that many housekeeping genes, including ribosomal RNAs and proteins, were present in these regions. These housekeeping genes are known to be highly transcribed throughout all stages of the bacterial life cycle. This finding led us to speculate that bacterial transcription may play a role in inhibiting transposase-mediated transposon insertion, especially in highly expressed genes. During transcription, the RNA polymerase holoenzyme, consisting of the RNA polymerase core enzyme and a sigma factor, interacts with promoter DNA to form a closed complex (Bae et al., 2015). This physical interaction effectively segregates the DNA, potentially limiting the access of transposase and impeding Tn5 insertion. Additionally, nucleoid-associated proteins modulate the compaction level of the bacterial chromosome through processes such as bending, wrapping, looping, and twisting DNA (Browning and Busby, 2016), thereby influencing its accessibility.

In a recent study on *Xanthomonas hortorum* pv. *vitians*, researchers identified a single essential gene encoding ribosephosphate pyrophosphokinase in the pentose phosphate pathway and another essential gene encoding phosphopyruvate hydratase in the glycolysis pathway (Morinière et al., 2021). However, our investigation did not identify any essential genes associated with these two pathways in Xcc CQ13. Instead, we found proteins related to the Entner-Doudoroff pathway, which is believed to play a significant role in the decomposition of most absorbed glucose in *Xanthomonas campestris* pv. *campestris* (Frese et al., 2014). These findings led us to conclude that in a nutrient-rich medium, no single pathway exhibited a clear advantage for the survival of Xcc CQ13. The significance of a particular pathway may vary depending on other factors in the natural environment, such as the availability of specific carbon substrates. Understanding these variations could be crucial in identifying pathway-specific drug targets, potentially contributing to the development of targeted therapies for controlling *Xanthomonas* infections.

## Experimental procedures

### Bacterial strains, plasmids, primers, and growth condition

The bacterial strains and plasmids used in this study are listed in Supplementary Table 7. The primers used in this study are listed in Supplementary Table 8. *Xanthomonas citri* strain CQ13 (Wu et al., 2022) was used to construct mutant libraries and was routinely cultured in Difco Nutrient Broth (NB) medium or on Difco Nutrient Agar (NA) at a temperature of 30 _. *E. coli* DH5α carrying either pLN2 or pLLN2 plasmid was cultured at 37 _ in Luria Broth medium. Kanamycin (50 μg ml^-1^) was supplemented when needed. Strains and mutant libraries were stored in the medium supplemented with 25% glycerol at -80□.

### Plasmid construction

The plasmid pLN2 was derived from the previous published pMCS2-transposase with minor modifications (Christen et al., 2011). The *lac* promoter was amplified from pBBR1MCS-2 and inserted into the upstream region of transposase gene. The PCR-amplified *lac* promoter fragment and the linearized pMCS2-transposase vector were assembled using Gibson assembly, resulting in the construction of pLN2. To address potential polar effects on downstream gene expression at the site of mutagenesis, plasmid pLLN2 was generated using pLN2 as the backbone. In this process, the second *lac* promoter was amplified and inserted outward-facingly at the end of the Tn5 transposon 3’ME (3’ mosaic end) sequence. This modification allowed for the creation of pLLN2, which could help minimize the potential impact on downstream gene expression caused by the transposon insertion.

### Construction of Tn5 transposon mutant library

A highly saturated transposon mutant library was generated using Tn5 transposon insertion, following a previously established protocol with certain modifications (Christen et al., 2011). The plasmids pLN2 and pLLN2 were introduced into the wild-type Xcc strain CQ13 by electroporation when the cells reached an optical density (OD600) of 0.4. After transformation, the Tn5 insertion mutants were subjected to recovery in NB medium supplemented with 1% sucrose for a duration of 3 hours. Subsequently, the recovered mutants were uniformly spread on NA plates containing kanamycin for selection. The resulting mutants from each library were harvested and pooled separately, designated as T1 and T2, respectively. These pooled mutants were pelleted through centrifugation at 10,000 g for 5 minutes at 4 °C and subsequently resuspended in NB medium supplemented with 20% (v/v) glycerol. The resuspended mutant pools were stored at -80 °C for long-term preservation.

### Library preparation for Illumina sequencing

The genomic DNA from the harvested pooled library was extracted using the CTAB method. For library preparation, restriction digestion and ligation steps were performed following the instructions provided by the manufacturer (New England Biolabs). The genomic DNA was digested using the restriction enzymes *Nco*□, *Sph*□, *Not*□, *Mfe*□, *Mlu*□ and *Bam*H□, respectively. Subsequently, the digested products underwent self-ligation using T4 DNA ligase at 16^°^C overnight. The reaction products were then ethanol-precipitated and washed twice with 70% (v/v) ethanol. The purified DNA was used as a template for reverse-amplification of the flanking genome sequences using an outward primer pair. The resulting PCR products were subjected to Illumina sequencing.

### Reads processing and identification of insertion sites

The data quality control was performed using FastQC v0.11.9. Reads with five consecutive base Ns or a length less than 20 were filtered out. Repeat sequences were filtered based on sequence using the "Rmdup" function of Seqkit v2.4.0. For genome mapping, the 25 bases at the 3’ end of the transposon sequence (GAGCTCACTTGTGTATAAGAGTCAG) were identified and extracted. The reads were then mapped onto the *Xanthomonas citri* CQ13 genome (accession number: CP096227). Comparisons were performed using the TPP, the original data processing software in Transit v3.2.7. The TPP software’s SAM format file output was converted to BAM using SAMtools v0.1.19, with unmatched sequences filtered out and soft-clipped reads removed. T1 and T2 insertion sites were combined, and repeated sites were deleted to obtain preliminary experimental results. The genome-wide distribution of insertion loci was visualized using the Lianchuan Biological Cloud Platform (https://www.omicstudio.cn/analysis) (Lyu et al., 2023). The insertion loci and GC skew were plotted using DNA Plotter. The correlation between transposon insertion abundance (the number of insertions in each gene vs. the number of insertions on the chromosome) of T1 and T2 was calculated using the R package "corrplot".

### Prediction of essential and high-fitness gene elements

In Transit v3.2.7, the Tn5Gaps tool was used to estimate gene essentiality. The replication mode was set as "Mean", and all other parameters were kept as default. The assessment was performed on the T1, T2, and fusion libraries, respectively. To determine the essential genes, *P*-values were calculated using a method described previously (Christen et al., 2011). Genes with a *P*-value smaller than 0.05 and fewer than 10 insertions were classified as essential genes. On the other hand, genes with a *P*-value smaller than 0.05 and more than 10 insertions were categorized as high-fitness cost genes.

To predict the origin of replication, the GC cumulative skew was calculated at each 200 bp interval of the genome using TBtools-II v1.113. The cumulative values of the GC skew were then computed using the *sum* function, and the position corresponding to the minimum value was identified as the predicted origin of replication. The MEME Suite (https://meme-suite.org/meme/) was used to explore the conserved motifs of the replication origin. Protein domain prediction was performed by SMART (https://smart.embl-heidelberg.de/).

### Gene enrichment and pathway mapping

The essential genes were assigned to Clusters of Orthologous Genes (COG) using eggnog-mapper v2 (http://eggnog-mapper.embl.de/). Fold enrichment of genes in a certain COG category was calculated as the ratio of the proportion of a given COG category in the essential gene set to its overall proportion in the entire genome and then log-transformed to determine overrepresented and underrepresented COG categories. The K number set based on KEGG (Kyoto Encyclopedia of Genes and Genomes) database converted from our essential gene set was retrieved from BLASTKOALA (https://www.kegg.jp/blastkoala/). Cellular pathways were generated by mapping K numbers to KEGG pathways with KEGG MAPPER (https://www.genome.jp/kegg/mapper/reconstruct.html) and gene enrichment analysis based on KEGG pathways was performed using R package clusterProfiler.

### Conservation and distribution of essential genes across the whole genome

To study the conservation of essential genes in Xcc CQ13 at the bacterial kingdom and genus levels, 66 essential gene sets from various bacteria were downloaded from the Database of Essential Genes (DEG15). For each gene set, BLASTP was employed with specific criteria: a minimum amino acid identity of 30% and an e-value threshold of less than 10^-5^. Homologues essential genes were searched using BLASTP with the same parameters as mentioned above. The phylogenetic tree based on the complete genomes of these 18 bacteria was constructed using BV-BRC (https://www.bv-brc.org/).

Sliding window analysis, with a window size of 500 kbp and a step size of 10 kbp, spanning the entire genomes of both bacterial species, was conducted to gain insights into the distribution and enrichment of essential genes in Xcc CQ13 and *E. coli*. For every step of the window, we counted the number of essential genes (n) and the total number of genes found within the window. In addition, we calculated the total number of essential genes and the total number of genes within the complete Xcc CQ13 and *E.coli* genome. *P*-values were calculated as the probability of detecting essential genes (^≥^n) within a given window by using a hypergeometric distribution. Finally, the *P*-value of every step of the moving window was collected and plotted as -log (*P*-value). The genome coordinates were calculated as the order of the window starting from the *dnaA* gene divided by the total number of windows collected in the whole genome.

## Acknowledgements

This study was supported by the Guangdong Basic and Applied Basic Research Foundation (2022A1515010905), the Open Project of BGI-Shenzhen (BGIRSZ20220003) and the Fundamental Research Funds for the Central Universities, Sun Yat-sen University (23qnpy41).

## Data availability statement

The Illumina sequencing raw data for this study can be accessed from the BioSample database under the following accession numbers: SAMN36786214, SAMN36786215, SAMN36786216, SAMN36786217, SAMN36786218, SAMN36786219, SAMN36786220, SAMN36786221, and SAMN36786222. All other data are available on request from the corresponding author.

## Supporting Information legends

**Supplementary Figure 1.** Multiple sequence alignment analysis of aspartate-semialdehyde dehydrogenase protein in *Xanthomonas* and homologs in other bacteria. The aspartate-semialdehyde dehydrogenase protein sequences from Xcc CQ13and its homologs in other bacteria were aligned using Blast and visualized with Jalview 2.11.2.71. Accession numbers for the homologs are as follows: *Pseudomonas* sp. HMWF010, PTT77345.1; *Escherichia coli*, WP_244563191.1; *Pseudoxanthomonas suwonensis*, PZO58311.1. The alignment displays conserved amino acids, indicated by their percentage identity in each column. The degree of identity is represented by color, with red indicating higher identity. Gaps in the alignment are denoted by "-".

**Supplementary Figure 2.** Prediction of the replication origin in Xcc CQ13. **(a)** The 273-bp genomic DNA sequence of the predicted origin of replication in Xcc CQ13 is displayed. The sequence containing the predicted conserved motifs is highlighted in red and bold, while the AT-rich sequence is underlined. These regions are proposed to be the putative binding sites for the replication initiator protein DnaA, critical for the initiation of DNA replication. **(b)** The consensus sequence of the predicted motifs is depicted, showing the relative size of the letters, which indicates their frequency in the sequence. The height of each letter is proportional to the frequency of occurrence of the corresponding base at that location.

## Notes

### Competing Interest Statement

The authors have declared no competing interest.

