## Supplemental Table 7 for "The essential genome of *Xanthomonas citri*"

**Supplementary Table 7. Strains and plasmids used in this study.**

| Name | Description | Source |
| --- | --- | --- |
| ***E. coli* strains** |  |  |
| NEB5alpha | general cloning strain | NEB |
| ***Xanthomonas* strains** |  |  |
| *Xanthomonas citri* subsp. *citri* CQ13 | The strain causes citrus canker disease | This study |
| **Plasmids** |  |  |
| pNPTS138 | suicide vector | Lab collection |
| pMCS2-transposase (pN2) | The vector used in *Caulobacter* | (Christen et al., 2011) |
| pLN2 | pN2 derivatives. The expression of transposase is driven by *lac* promoter. | This study |
| pLLN2 | pLN2 derivatives. An additional *lac* promoter is inserted outward-facingly at the end of the Tn5 transposon | This study |

Reference：

Christen, B., Abeliuk, E., Collier, J.M., Kalogeraki, V.S., Passarelli, B., Coller, J.A., et al. (2011) The essential genome of a bacterium. *Molecular Systems Biology*, 7, 528. https://doi.org/10.1038/msb.2011.58.
