## Supplemental Table 8 for "The essential genome of *Xanthomonas citri*"

**Supplementary Table 8. Primers used in this study.**

| **Primers for construction of plasmids** | |
| --- | --- |
| #1 | atggtaacgttcatgataacttc |
| #2 | ttccgttcaggacgctacttg |
| #3 | aagtagcgtcctgaacggaagtgatggcttccatgtcggcag |
| #4 | gttatcatgaacgttaccatagctgtttcctgtgtgaaattgttatc |
| #5 | tcgcgagacgtccaattgcagcccaatacgcaaaccgcctctc |
| #6 | ctcttgatcagatctggtacagctgtttcctgtgtgaaattgttatc |
| **Primers for amplification of genomic DNA** | |
| #7 | ctttgagtgagctgataccgc |
| #8 | cgcgagacgtccaattgcatat |
| #9 | gcctttgagtgagctgataccgc |
| #10 | tgtttgtcgcattatacgcaaggc |
| #11 | gcaagacgtttcccgttgaatatgg |
| #12 | cagataaaacgaaaggcccagtctttc |
