## Supplementary figures and images for "The essential genome of *Xanthomonas citri*"

### Supplemental Figure 1

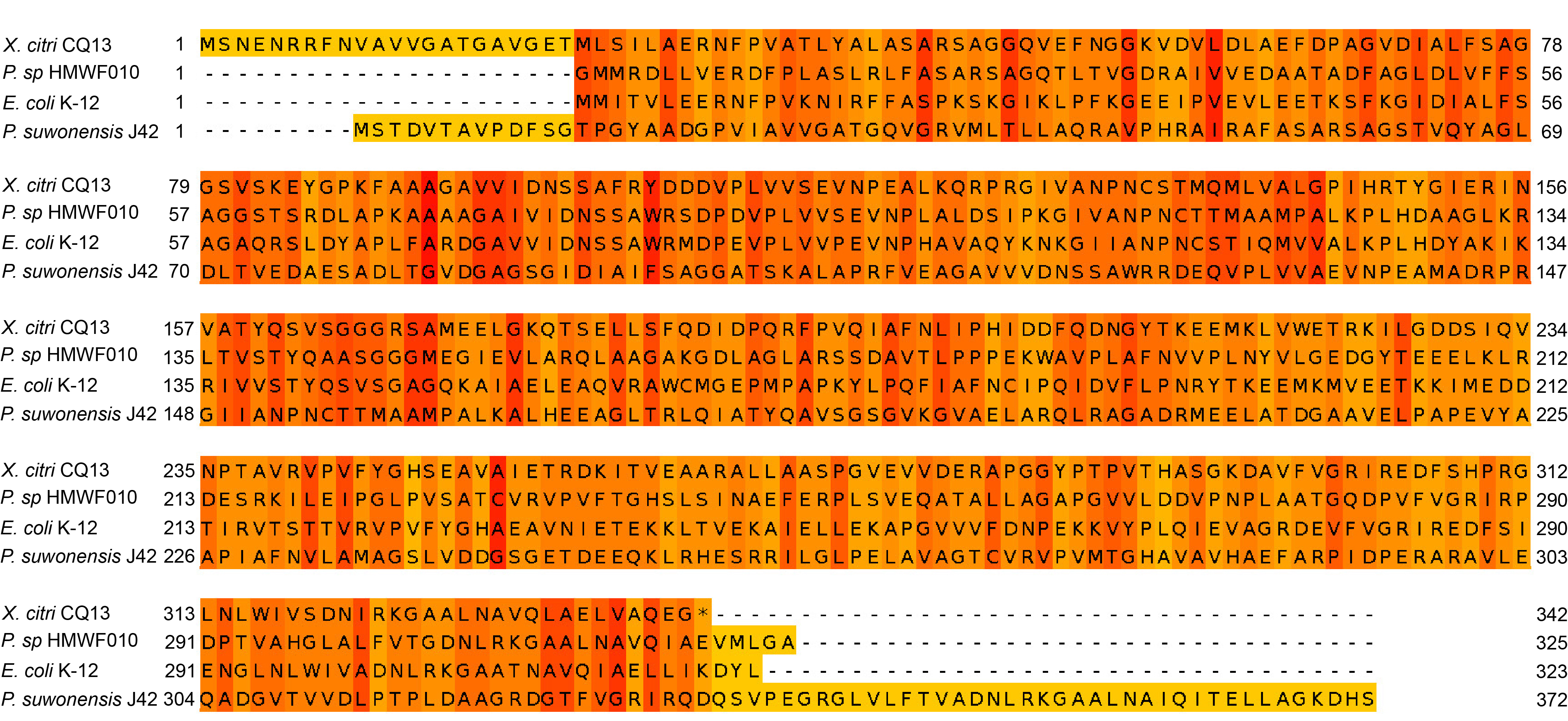

### Supplemental Figure 2

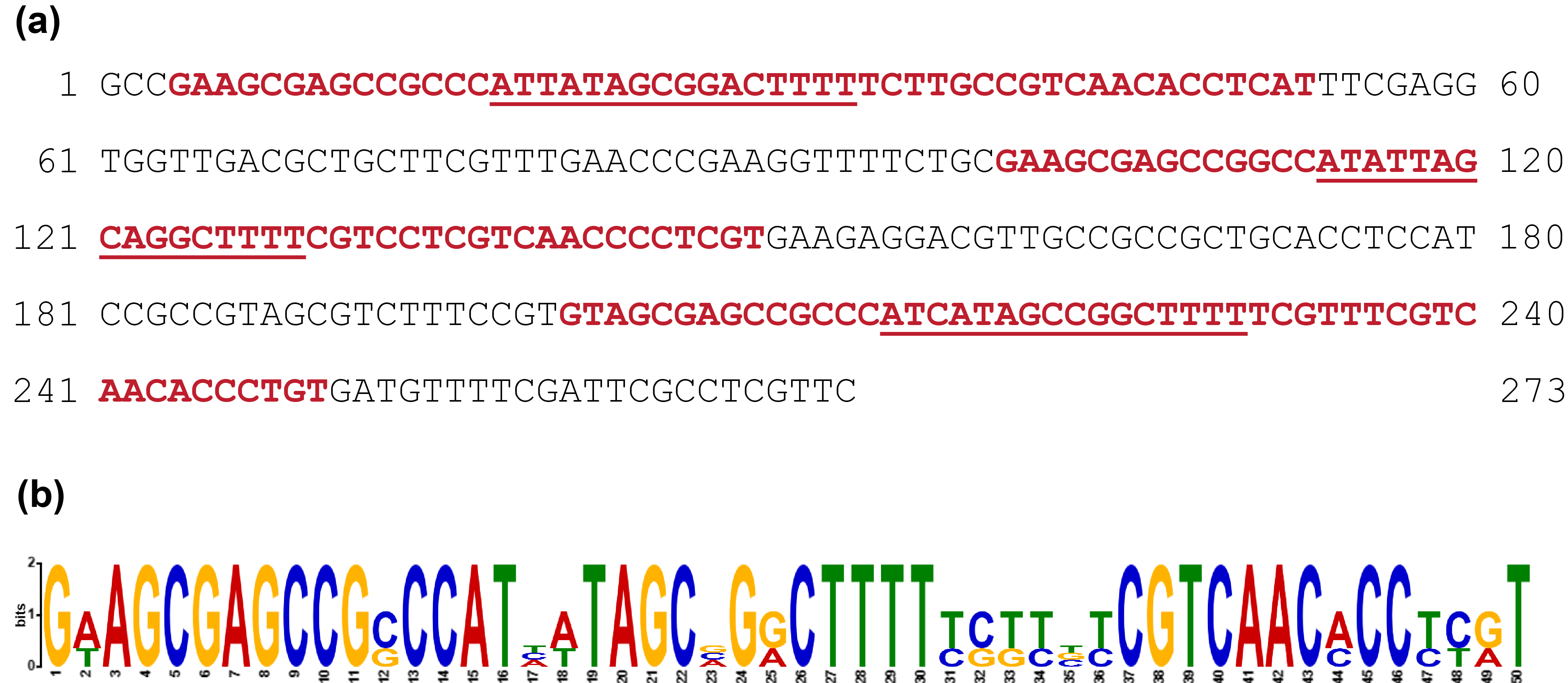
